# Hypothalamus amyloid levels are associated with early sex-dependent alterations in energy homeostasis in TgF344-AD rats

**DOI:** 10.64898/2026.03.08.710398

**Authors:** Caleb M. Levine, Cameron Caggiano, Thea Anderson, Michael A Kelberman, David Weinshenker, Hannah L Lail, Desiree Wanders, Debra A Bangasser, Scott E Kanoski, Marise B. Parent

## Abstract

We reported previously that diet-induced obesity exacerbates early-stage Alzheimer’s disease (AD)-like pathology in TgF344-AD rats. Our findings also suggested that TgF344-AD rats may be prone to weight gain during early AD development, which we assessed here. Energy intake, body composition, and the impact of glucose administration on blood glucose were also assessed. Body temperature, intrascapular brown adipose tissue (iBAT) mass, and iBAT uncoupling protein-1 (UCP1) expression were used as indicators of thermogenic function. Soluble amyloid β_40_ (Aβ_40_) and Aβ_42_ were quantified in hypothalamus. Male TgF344-AD rats began to outweigh wildtype (WT) littermates by 5 weeks of age; this increase emerged later in female TgF344-AD rats (~5 months). Female TgF344-AD rats ingested more energy from chow and a high fat, high sugar (HFHS) diet, gained more weight on the HFHS diet, and had lower UCP1 than WT rats, effects not observed in male TgF344-AD rats. Surprisingly, male and female TgF344-AD rats had increased body temperatures. This was restricted to the dark phase in females, which is when they ingest excess calories. Finally, the HFHS diet disrupted glucose regulation in male but not female TgF344-AD rats. These findings suggest that increases in energy intake and decreases in UCP1 may contribute to the additional weight gain in female TgF344-AD rats. The causes for these increases in males remain unclear. Hypothalamic Aβ_42_ correlated with glucose dysregulation in male TgF344-AD rats and BAT mass in female TgF344-AD rats, raising the possibility that increases in Aβ_42_ in hypothalamus produce sex-specific disruptions in energy homeostasis.

## 1. INTRODUCTION

Alzheimer’s disease (AD) is the most common form of dementia, affecting an estimated 416 million people worldwide, and AD prevalence is expected to continue to increase [1,2]. Currently, the most effective treatments available only modestly attenuate AD symptoms and slow the rate of decline, and neither curative nor preventative solutions are available [3,4]. AD is characterized by the progressive deposition of amyloid-β peptides (Aβ) as plaques, the development of neurofibrillary tangles (NFTs) composed primarily of the hyperphosphorylated microtubule-associated protein tau (ptau), and eventual neuronal death [5,6]. AD progresses in stages, starting with decades of preclinical asymptomatic neuropathology that typically starts in middle age. This pathology includes elevated Aβ and changes in tau that often go undetected until mild cognitive impairment (MCI) appears during the prodromal phase, which in turn lasts until frank dementia emerges during the clinical phase [7-12].

Many risk factors associated with AD exert their effects during the preclinical and prodromal periods (i.e., during middle age). Of note, midlife obesity doubles AD risk and predicts earlier AD onset and more severe pathology [13-15]. This is a significant concern because the worldwide prevalence of obesity has nearly tripled since 1975 [16]. Although midlife obesity is associated with increased AD risk [17-22], late-life obesity is not, suggesting that the risk interactions between obesity and AD occur during preclinical/prodromal AD stages. Even though revised guidelines have allowed the diagnosis of AD at earlier stages of pathology based on positive imaging or blood biomarkers, not all will go on to develop symptoms of AD [23]. As a result, it remains very challenging to investigate the mechanisms underlying risk factors during preclinical and prodromal AD.

Given these challenges, nonhuman animal models are vital for investigating the neural changes that occur during early AD development and for understanding how risk factors increase the likelihood of transitioning from early to later stages. Of all rodent models currently available, the TgF344-AD rat, which expresses the Swedish mutant human amyloid precursor protein (APPsw) and Δexon9 mutant human presenilin-1 (PS1 ΔE9), appears to most closely recapitulate human AD trajectory [24]. TgF344-AD rats develop elevations in Aβ oligomers (Aβos), intraneuronal Aβ, plaques, progressive memory deficits, neuroinflammation, gliosis, and apoptotic neuronal injury/loss in an age-dependent manner [24,25]. Unique to this model, Aβ triggers endogenous tauopathy consisting of NFT-like aggregated rat ptau in brain areas relevant to early AD pathology, such as the locus coeruleus, hippocampus, cortex, and entorhinal cortex [24-29]. This is a significant advantage of this model because tau pathology is more closely related to the etiology of neurodegeneration and dementia than Aβ [30,31]. Importantly, neuropathology in TgF344-AD rats proceeds through characteristic preclinical, prodromal, and “clinical” stages [24,25,28,32]. The prodromal period appears to begin between 4-5 months of age, with increased anxiety, subtle learning deficits, and impaired functional connectivity [33,34]. At 6 months, there are significant elevations in soluble Aβos, neuroinflammation, and mild cognitive impairment [24,25]. At 9 months, there is widespread amyloid plaque deposition, tau pathology, and neurodegeneration [26,35]. The “clinical” period in TgF344-AD rats emerges at approximately 14-16 months and is characterized by significant cognitive deficits, aggregated ptau, and neuronal loss [24,28,36-38]. The protracted prodromal stage in TgF344-AD rats relative to other rodent AD models provides the opportunity to test the neurobiological mechanisms by which risk factors during this stage exacerbate later AD pathology, employing study designs that are not possible in human subjects.

Evidence suggests that energy homeostasis may be disrupted in TgF344-AD rats. For example, TgF344-AD rats fed standard chow weigh more than their WT counterparts [37,39,40] and have elevated blood lipid levels [40]. Moreover, feeding high fat/high sugar (HFHS) diets to TgF344-AD rats, particularly females, produces larger elevations in weight gain and adiposity than in WT rats (Anderson et al., 2023; Almanza et al., 2024). The findings from the last two studies are particularly intriguing as they examined the effects of diet-induced weight gain during the prodromal period in these rats, when obesity is most impactful as a risk factor for future AD development in humans. This evidence of disrupted energy homeostasis during early AD development raises the possibility that the hypothalamus, which is critical for energy homeostasis [41,42], is impaired during early AD development in these rats. In support, Aβ plaques and NFTs have been observed in the hypothalamus of patients with AD, although most of this pathology has been identified in later stages of AD development [43]. Importantly, a recent systematic review demonstrated that patients with mild to moderate AD have reduced hypothalamic volume, suggesting that the hypothalamus may indeed be impacted early in AD [44]. Given the protracted time course of pathological changes that occur over decades in AD, these findings suggest that the hypothalamus may be implicated in AD prior to clinical diagnosis.

Based on the evidence reviewed above, the goal of the present study was to characterize energy homeostasis and peripheral metabolic function in TgF344-AD rats during the earliest stages of AD, as well as to determine whether soluble amyloid pathology is present in the hypothalamus during this period. If the findings indicated that the brain Aβ in these rats is associated with a propensity toward weight gain and metabolic disturbances, then this might suggest that disruptions in energy homeostasis are an early symptom of AD development that may potentially contribute to the risk effects of obesity during prodromal AD.

## 2. MATERIALS AND METHODS

### 2.1. Animals

A breeding colony was established at Georgia State University using hemizygous TgF344-AD sires and Fischer 344 WT dams (Charles River Laboratories, Kingston, NY). Pups were genotyped using ear tissue punch biopsies taken at weaning on postnatal (P) days 21-25 (Transnetyx, Memphis, TN). Unless otherwise stated, rats were housed in pairs or triads in ventilated polycarbonate cages in a temperature and humidity-controlled facility (25 °C, 40% humidity) on a 12 h light-dark cycle with lights on at 7 am. The rats were given *ad libitum* access to food and water. All animal procedures were approved by the Georgia State University Institutional Animal Care and Use Committee (IACUC), were carried out in accordance with NIH (National Research Council) Guide for the Care and Use of Laboratory Animals, and comply with ARRIVE (Animal Research: Reporting of In Vivo Experiments) guidelines.

### 2.2. Body mass

To determine when differences in body mass emerge, male and female offspring of each genotype were weighed weekly from weaning on P21 until P70. An independent cross-sectional assessment of body mass in male and female TgF344-AD and WT rats of different age groups was also obtained from a colony at Emory University. Those procedures were approved by the Emory University IACUC.

### 2.3. Energy intake

To assess energy intake, rats were housed individually at 4-5 months of age in cages equipped with food hoppers hanging from scales (TSE LabMaster Metabolic Research Platform; TSE Systems International, Chesterfield, Missouri). They were given free access to standard chow (LabDiet 5001; Richmond, IN), water, and a Nylabone for enrichment. The rats were given 3 days to acclimate to the new housing, and then chow intake was assessed for 4 days. On the 8^th^ day, all rats were given a palatable high calorie (kcal) HFHS diet that supplied 45% of the kcal from fat and half of the carbohydrates from sucrose (S. Table 1; Research Diets D12451; New Brunswick, NJ). Intake was assessed for 4 days yielding within-subject responses to both chow and the HFHS diet. Food intake was recorded every 27 min. Each morning, the recording was stopped at 10:00 a.m. for 30 min while rats were weighed; bedding was changed as necessary, and food hoppers/water bottles were refilled. Grams of food consumed were converted to kcal consumed based on caloric content of the diet (S. Table 1). To control for the effects of differences in body mass on intake, daily energy intake was normalized to the body mass recorded at the start of that day. Energy intake was analyzed separately during the light and dark phase of the light cycle.

### 2.4. Body temperature

Unpublished body temperature data obtained from rats used in the Anderson et al. (2023) study were analyzed to determine whether reductions in energy expenditure could account for the increases in body mass [45]. At 4 months of age, rats were surgically implanted with iButton temperature sensors (ThermoChron, Sidney, Australia) into the intraperitoneal cavity under deep isoflurane anesthesia (5% for initiation; 2% for maintenance; O_2_ flow rate: 1.5 L/min). The rats were given carprofen (5 mg/kg SC) prior to surgery and 24 h post-surgery. The iButtons were programmed to record temperatures starting ~2 weeks after implantation at 30-min increments with.0625°C precision, encased in histology grade paraffin wax, and sterilized with ethylene oxide gas. After 2 weeks of recovery, half of the rats in each group were randomly selected and placed on the HFHS diet. After 11 weeks of recording, the rats were euthanized as described in Anderson et al. (2023), the iButtons were retrieved, and the temperature data were extracted [45]. Temperature readings were binned by 30 min and aligned within ±12 min to the zeitgebers of lights-on and lights-off (e.g., the 7 am timepoint represents 6:48 am–7:12 am). The average measurement for each timepoint was taken across 8 weeks of recording.

### 2.5. Western blot analysis of iBAT

Western blot (Bio-Rad) was used to measure uncoupling protein 1 (UCP1) protein expression in interscapular brown adipose tissue (iBAT) tissue collected from the rats in our previous study [45]. Total lane protein (Sigma-Aldrich T54801) was used to normalize the results for statistical analyses. Western blots followed standard protocols as previously described [46]. Protein was extracted from iBAT using radioimmunoprecipitation assay lysis buffer (RIPA) containing a protease inhibitor cocktail (Sigma-Aldrich P8340) and phosphatase inhibitors (50 mM sodium fluoride, 1 mM sodium orthovanadate, 2.5 mM sodium pyrophosphate decahydrate, 10 mM 2,3-bisphosphoglyceric acid). Total protein concentration was determined with a detergent compatible protein assay and a microplate reader (Synergy HT, BioTek). Proteins were separated by 10% SDS-PAGE electrophoresis and transferred to a polyvinylidene difluoride membrane (Bio-Rad 10026933). Membranes were blocked in 5% non-fat dry milk for 1 h at room temperature followed by overnight incubation at 4°C in a primary antibody raised against the peptide sequence corresponding to amino acids 145 to 159 in UCP1 [47]. Membranes were then washed and incubated with secondary antibodies (Rabbit-CS7074S) at room temperature for 1 h. Protein expression was visualized using the Bio-Rad ChemiDoc Imaging System.

### 2.6. EchoMRI

A separate group of rats (~4.5 months old) were matched for body weight within each genotype and sex and assigned to either continue standard chow (S. Table 1; LabDiet 5001; Richmond, IN) or were switched to the HFHS diet (S. Table 1; Research Diets D12451; New Brunswick, NJ) and weighed weekly. After 8 weeks on diet, body composition was assessed using an EchoMRI-700 Whole Body Analyzer (EchoMRI™, Houston, TX; Figure 3A). Unanesthetized rats were guided into a ventilated plastic tube that was then inserted into the machine. Fat mass and lean body mass were averaged from duplicate 3-min readings. Measurements were calibrated to two known standard volumes of vegetable oil that were established through triplicate 3-min readings.

### 2.7. Glucose tolerance test

Fasting glucose tolerance was assessed in a random subset of these rats after 9 weeks on the diet. Food was removed between 1 to 3 h after the start of the light cycle, and rats were brought into the testing room to acclimate for 8 h +/-30 min prior to the first blood glucose collection. A small puncture was made on the tip of the tail using a disposable 28-gauge lancet (CVS Thin Lancets; CVS 235157) and a drop of blood was collected to determine baseline blood glucose levels using a glucose meter (CVS Advanced Glucose Meter 968874). Then, rats were given an injection of glucose (1.7 g/kg lean mass; IP) and returned to their home cages. Lean body mass from the EchoMRI conducted 1 week earlier was used to calculate dosing. Blood samples were collected 5, 10, 30, 45, 60, 90, and 120 min following the injection by milking the tail, forcing blood to displace the scab on the tail puncture. The first drop of blood was removed with a tissue, and the subsequent drop was sampled.

### 2.8. Tissue collection

After 11 weeks on their diets (7-7.5 months of age), rats were euthanized with sodium pentobarbital (Euthasol; 150 mg/kg IP; Virbac VINV-CIII-0001). The distance from the nose to the anus was measured to determine body length. iBAT, liver, heart, kidneys, and adrenal glands were harvested and weighed. Brains were extracted from the skull, the hypothalamus was gross-dissected, and then frozen on dry-ice and stored at −80°C.

### 2.9. ELISA analysis of soluble Aβ

The hypothalamic samples were suspended in 20 volumes of cell lysis buffer (10× Complete cell lysis buffer diluted to 1×; Cell Signaling Technology 9803) with 1% protease and phosphatase inhibitor cocktail (HALT; Thermofisher 78440), then homogenized with 20 strokes of a mortar and pestle on wet ice. After 30 min, the lysate was centrifuged at 16000 ×g for 30 min at 4 °C. The supernatant was then removed and frozen on dry ice until further analyses. Protein content was quantified by bicinchoninic acid assay (Pierce BCA; Thermofisher 23225) and then Aβ species were quantified using commercial ELISA kits selective for human Aβ42 (Thermofisher KHB3441) and human Aβ40 (Thermofisher KHB3481) following the manufacturer’s instructions. The samples were quantified using a four-parameter logistical regression (myassays.com) and normalized to the total protein concentration of the sample.

### 2.10. Statistics

Experimenters were blinded to the condition of the animals and samples during data collection for *in vivo* and post-mortem assessments. The data were tested for normality, homogeneity of variance, and covariance using R. For data that were normally distributed, values exceeding 2.5 standard deviations from the mean were identified as outliers and excluded from subsequent statistical analyses (S. Table 2). For data that were not normally distributed, values exceeding 2.5 times the interquartile range (IQR) were identified as outliers. Since litter size influences a variety of metabolic and morphometric measures [48,49], a mixed linear model (Kenward–Roger approximation) with litter size included as a random effect was used to analyze intake with sex, diet, genotype, and time as factors (S. Figure 1A). When there were main effects of sex or interactions between sex and genotype, diet or time, a subsequent mixed linear model was conducted separately in males and females with only genotype and diet (or genotype, diet, and time) as fixed effects. Effect sizes are presented as partial eta squared 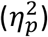. Pairwise t-tests were conducted on estimated marginal means using Bonferroni correction. Total area under the curve (tAUC) was calculated for the length of the glucose tolerance test (GTT) using the trapezoidal rule [50]. The GTT was then broken down into 30-minute segments, and an AUC was taken for each segment (0-30 min; 30-60 min; 60-90 min; 90-120 min). Significant interactions between genotype and diet within each sex were analyzed with Bonferroni-corrected pairwise t-tests. The Aβ ELISA data were analyzed using a 2-way ANOVA (sex x diet). Pearson’s correlations were computed between hypothalamic Aβ_42_ or Aβ_40_ and iBAT mass or tAUC and between iBAT UCP1 and terminal body mass. A probability of *p* <.05 was considered statistically significant. Averages ± standard error of the mean (SEM) are plotted for the depicted groups.

## 3. RESULTS

### 3.1. Male and female TgF344-AD rats weigh more than WT rats across several ages during the prodromal period

Rats were weighed weekly starting after weaning (P21-P24) to identify when the weight of TgF344-AD rats begins to diverge from their WT counterparts (Figure 1A, B). Female TgF344-AD rats did not significantly differ from WT female rats during this period and planned post-hoc comparisons failed to identify any genotypic effects (Figure 1A). For males, there was a significant interaction between genotype and age (F_1,567_ = 19.79, p = 9.68×10^−6^, 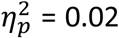; Figure 1B), such that male TgF344-AD rats had higher body mass than male WT rats (F_1,567_ = 56.17, p = 2.56×10^−13^, 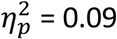). Planned post-hoc comparisons showed that by week 5, male TgF344-AD rats weighed significantly more than male WT rats (t_560.5_ = 2.845, p =.032).

**Figure 1.**
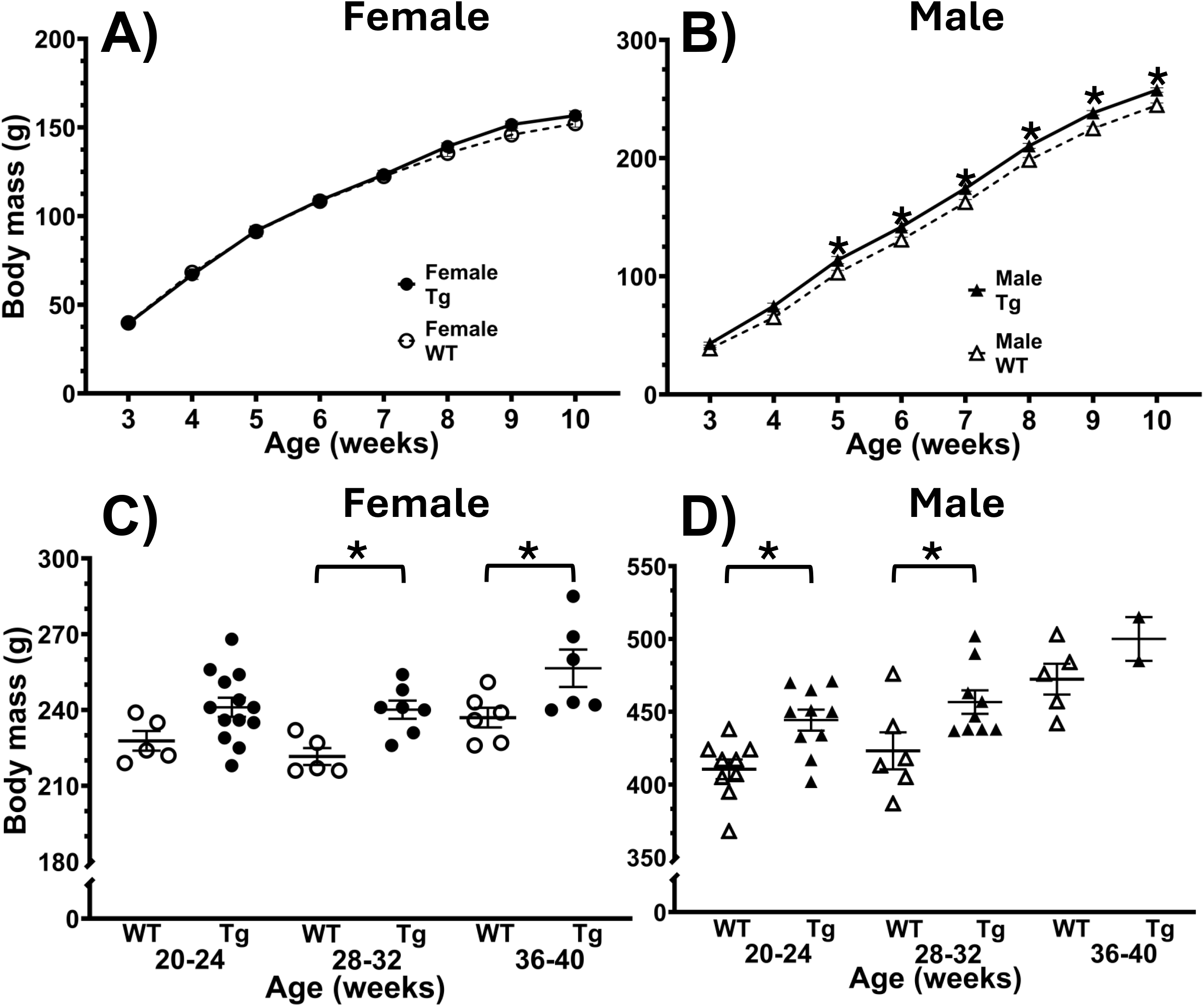
Male and female TgF344-AD rats weigh more than WT controls throughout most of the prodromal period. Male TgF344-AD rats begin to weigh more than WT controls during adolescence. **A)** Female TgF344-AD rats did not display any differences during the first 10 weeks of life (n = 23-38/gp). **B)** Male TgF344-AD rats began to weigh more than WT controls at 5 weeks of age (n = 23-38/gp). **C)** Female TgF344-AD rats weighed significantly more than WT littermate controls at 28-32 and 36-40 weeks of age (n = 5-13/gp). **D)** Male TgF344-AD rats weighed significantly more than WT littermate controls at 20-24 and 28-32 weeks of age (n = 2-12/gp). *WT = Wild-type, Tg = TgF344-AD*, ^*\**^ *= p <0.05*

A cross-sectional analysis of body mass from a different cohort of TgF344-AD rats at multiple ages showed that female and male TgF344-AD rats weighed more than their WT counterparts (female: F_1,36_ = 18.377, p =.000129, 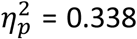; Figure 1C; male: F_1,34_ = 12.44, p =.0012, 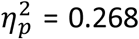; Figure 1D). Planned post-hoc comparisons indicated that female TgF344-AD rats began to weigh significantly more than female WT rats at 28-32 weeks of age (t = 2.58, df = 34, p =.042); meanwhile, the male TgF344-AD rats already weighed significantly more than male WT rats by20-24 weeks of age (t = 2.59, df = 36, p =.042).

### 3.2. Female TgF344-AD rats eat more than WT females

Energy intake was assessed in the rats from Figure 1 when they were 4.5-5 months of age (Figure 2A). The number of kcals consumed were divided by body mass to control for the effects of body mass on energy intake. There was a significant interaction between genotype and light cycle for energy intake (F_1,229.6_ = 11.09, p =.001, 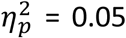) such that female TgF344-AD rats consumed more calories than female WT rats during the dark cycle (F_1,93.199_ = 32.854, p = 1.215×10^−7^, 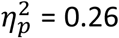; Figure 2B). This effect in females was accompanied by an interaction with diet (F_1,93.001_ = 4.754, p =.0318, 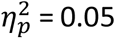), such that there was a greater mean difference in intake between WT and TgF344-AD rats on the HFHS diet (t_229.91_ = 3.98, p = 9.24×10^−5^) than on the chow diet. Female TgF344-AD rats did not have elevated intake during the light phase (Figure 2C), and intake did not differ between male WT and TgF344-AD rats during either the light or dark portion of the cycle (Figure 2D, E).

**Figure 2.**
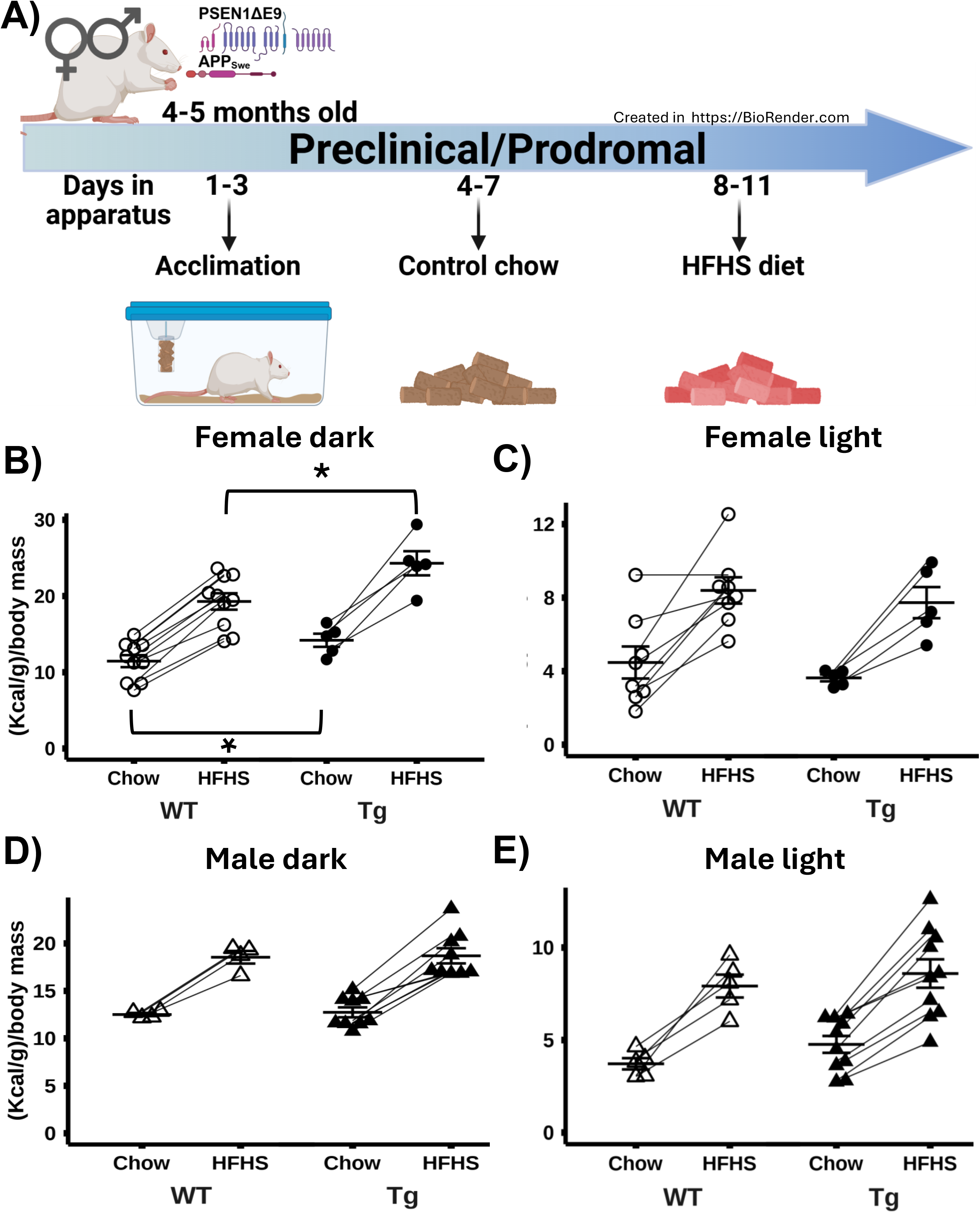
Female TgF344-AD rats consume more calories than WT female rats during the dark phase of the light cycle. **A)** Experimental timeline. **B)** During the dark phase, female TgF344-AD rats consumed more of the control and HFHS diet than their WT counterparts (WT: n = 10, Tg: n = 5). **C)** There were no significant effects of genotype on intake during the light phase in females (WT: n = 8, Tg: n = 5). **D)** There were no significant effects of genotype on intake during the dark (WT: n = 4, Tg: n = 9) or **E)** light in male rats (WT: n = 5, Tg: n = 10). *WT = Wild-type, Tg = TgF344-AD, HFHS = High-fat high-sugar diet*, ^*\**^ *= p <0.05*

### 3.3. Chronic consumption of a HFHS diet produces bigger increases in body weight in female TgF344-AD rats than WT rats

A different group of rats were placed on the HFHS diet for 11 weeks to assess the impact of chronic consumption of the diet (Figure 3A). There was a significant main effect of genotype in female rats (F_1,146.5_ = 6.11, p = 1.46×10^−2^, 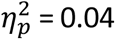; Figure 3B), with TgF344-AD rats weighing more regardless of diet. Genotype did not affect body mass in male rats during the study period (Figure 3C). Female (t = 2.47, p =.015), but not male TgF344-AD rats weighed more than WT rats at the time of euthanasia (Figure 3D, E). The higher body mass in female rats was not due to increased body size (S. Figure 2A).

**Figure 3.**
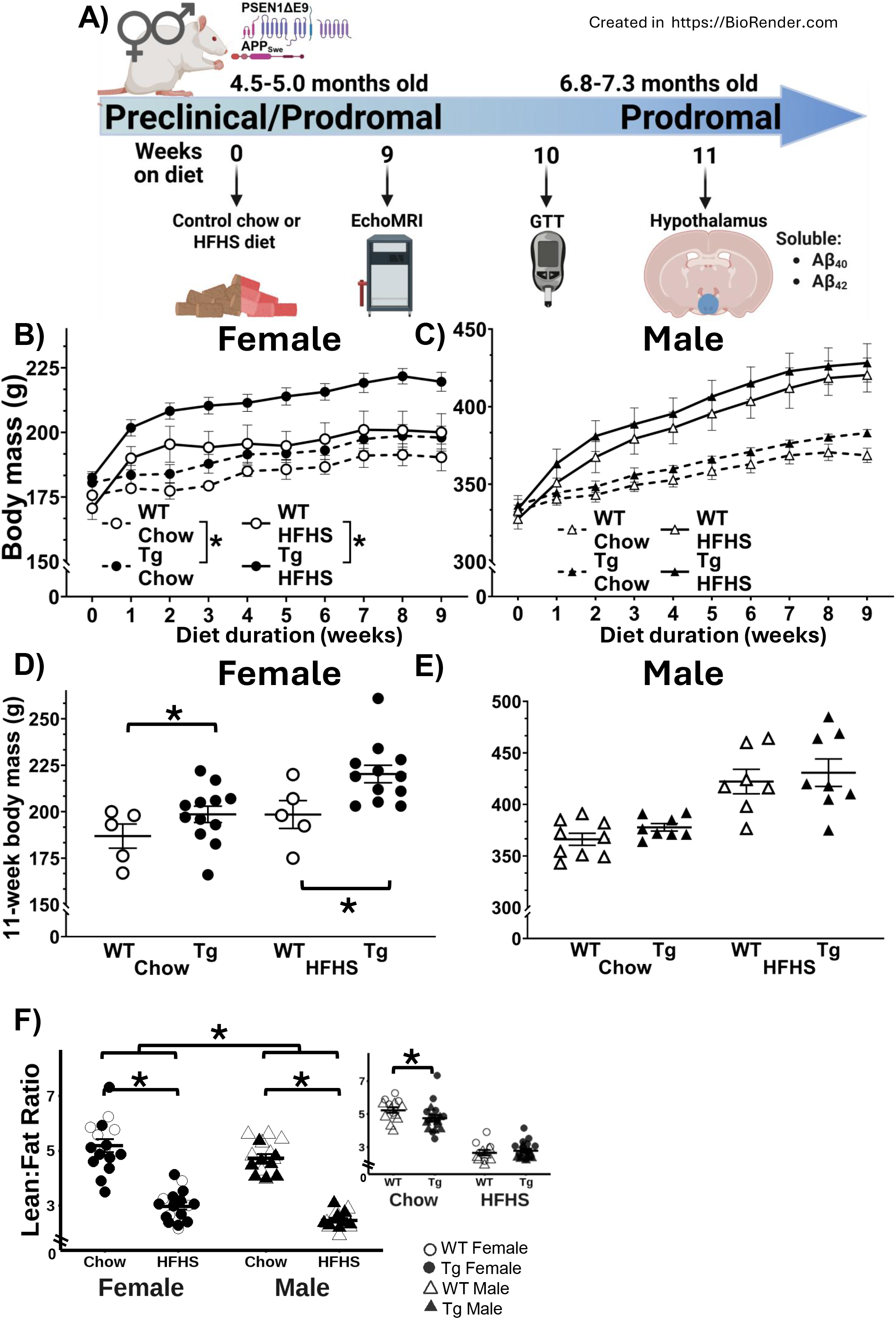
Female TgF344-AD rats accrue more body mass than WT females, especially on the HFHS diet. **A)** Experimental timeline. **B)** Body mass was elevated in female TgF344-AD rats fed the chow or HFHS diet (WT-Chow: n = 5, WT-HFHS: n = 5, Tg-Chow: n = 11, Tg HFHS: n = 11). **C)** This effect was not seen in male TgF344-AD rats (WT-Chow: n = 9, WT-HFHS: n = 7, Tg-Chow: n = 7, Tg HFHS: n = 8). **D)** Female TgF344-AD rats weighed more than WT rats at the end of the experiment (WT-Chow: n = 5, WT-HFHS: n = 5, Tg-Chow: n = 10, Tg HFHS: n = 11), **E)** but male TgF344-AD rats did not (WT-Chow: n = 9, WT-HFHS: n = 7, Tg-Chow: n = 8, Tg HFHS: n = 8). **F)** Chow-fed TgF344-AD rats had a lower lean-to-fat mass ratio than their WT counterparts regardless of sex (Male: WT-Chow: n = 9, WT-HFHS: n = 7, Tg-Chow: n = 8, Tg-HFHS: n = 8; Female: WT-Chow: n = 5, WT-HFHS: n = 5, Tg-Chow: n = 11, Tg-HFHS: n = 12). *WT = Wild-type, Tg = TgF344-AD, HFHS = High-fat high-sugar diet*, ^*\**^ *= p <0.05*

### 3.4. Chow-fed TgF344-AD rats have a reduced lean-to-fat mass ratio regardless of sex

Quantitative magnetic resonance results showed that diet and genotype interacted (F_1,56.01_= 4.22, p =.0447, 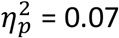) so that chow (t = 2.74, df = 53.47, p = 0.0333), but not HFHS diet-fed (t =.238, df = 56.95, p = 0.81) TgF344-AD rats had a lower lean-to-fat ratio than their WT counterparts regardless of sex. Irrespective of genotype, female rats had a higher lean-to-fat ratio than male rats (F_1,56.28_ = 9.75, p =.003, 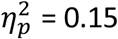; Figure 3F), and that the HFHS diet reduced the ratio of lean-to-fat in both sexes (Male: F_1,18.59_ = 2.56, p = 2.56×10^−10^, 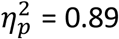; Female: F_1,28.17_ = 69.03, p = 4.64×10^−9^, 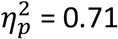).

Regardless of sex, HFHS-fed rats had increased percent body fat (F_1,56.79_ = 237.31, p = 4.84×10^−22^, 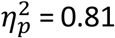; S. Figure 1B) but there was no effect of genotype on adiposity (F_1,56.80_ = 0.71, p =.403, 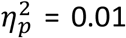). HFHS-fed rats also had reduced percent lean mass (F_1,54.26_ = 156.92, p = 3.70×10^−18^, 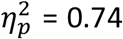; S. Figure 1C). The impact of diet on percent lean mass depended on sex (F_1,56_ = 6.42, p =.014, 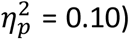 Specifically, post-hoc analyses showed that diet and genotype interacted in males (F_1,23.16_ = 7.41, p =.012, 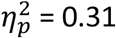), but not females (F_1,28.46_ = 0.84, p =.368, 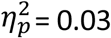) such that chow-fed TgF344-AD males had less percent lean mass than chow-fed WT males (t = 3.03, df = 22.15, p =.017).

### 3.5. Male TgF344-AD rats fed the HFHS diet have impaired glucose regulation

Glucose administration significantly increased blood glucose concentrations in all groups (F_1,486.41_ = 17.46, p = 1.12×10^−20^, 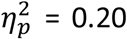; Figure 4A, B). There was a significant interaction between genotype and diet (F_1,439.78_ = 5.78, p =.017, 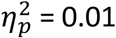) that was specific to males (F_1,221.85_ = 5.47, p =.02, 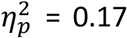; females F_1,193.05_ = 1.14, p =.287, 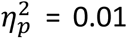). Specifically, glucose administration produced bigger increases in blood glucose concentrations in HFHS-fed male TgF344-AD than WT males fed the same diet (t = 3.57, df = 230.53, p =.002; Figure 4B). When the AUC was broken down into 30-min segments, there was an interaction between sex and genotype from 0-30 min (F_1,31.8_ = 4.61, p =.039, 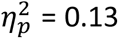; Figure 4C), and main effects of sex from 30-60 min (F_1,28.85_ = 5.47, p =.026, 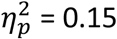; Figure 4D), 60-90 min (F_1, 20_ = 17.89, p = 4.11 × 10^−4^, 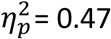; Figure 4E), and 90-120 min (F_1,20_ = 14.73, p =.001, 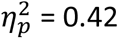; Figure 4F). During the post-injection incline from 0-30 min (Figure 4C), diet and genotype interacted in male rats (F_1,16.31_ = 5.32, p =.035, 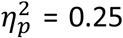). Specifically, HFHS-fed TgF344-AD rats had a greater AUC than their HFHS-fed WT counterparts (t = 3.51, df = 16.23, p =.011) and chow-fed TgF344-AD rats (t = 2.92, df = 16.52, p =.039). As blood sugar levels declined from 30-60 min (Figure 4D), diet and genotype interacted in male rats (F_1,15.58_ = 4.69, p =.046, 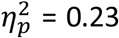), such that HFHS-fed male TgF344-AD rats had a greater AUC than WT rats fed the same diet (t = 3.44, df = 15.58, p =.014). HFHS-fed male rats had a greater AUC than their chow-fed counterparts as blood glucose levels continued to decline between 60-90 min (F_1,9.91_ = 7.16, p =.023, 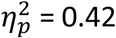; Figure 4E) and 90-120 min (F_1,10.16_ = 7.68, p =.019, 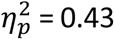; Figure 4F). TgF344-AD males had a greater tAUC than WT males (F_1,15.68_ = 4.67, p =.047, 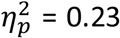; Figure 4G), but genotype did not interact with HFHS diet to further increase AUC (F_1,15.68_ = 2.72, p =.119, 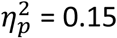).

**Figure 4:**
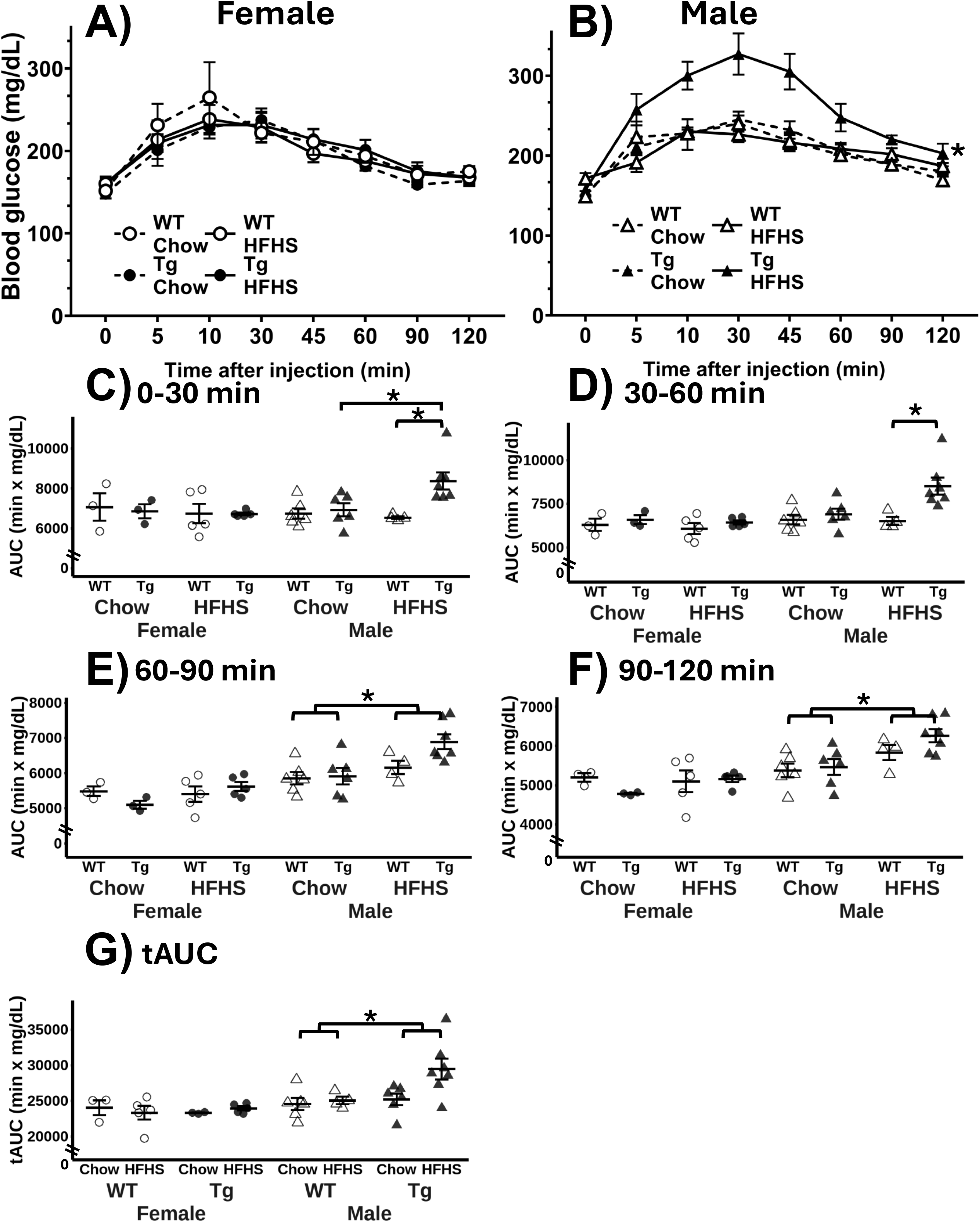
Male but not female TgF344-AD rats have disrupted glucose homeostasis. **A)** Female rats (6.3-7 months) did not exhibit diet- or genotype-induced disruptions in glucose regulation (WT-Chow: n = 3, WT-HFHS: n = 5, Tg-Chow: n = 4, Tg-HFHS: n = 5). **B)** Glucose administration increased blood glucose levels more in male TgF344-AD rats fed the HFHS diet than any other group of male rats (WT-Chow: n = 6, WT-HFHS: n = 4, Tg-Chow: n = 6, Tg-HFHS: n = 7). **C-F)** The HFHS diet increased 30 minute AUC in male, but not female TgF344-AD rats during the GTT. **G)** Male TgF344-AD rats had elevated tAUC. (Male: WT-Chow: n = 6, WT-HFHS: n = 4, Tg-Chow: n = 6, Tg-HFHS: n = 7; Female: WT-Chow: n = 3, WT-HFHS: n = 5, Tg-Chow: n = 3, Tg-HFHS: n = 5) *WT = Wild-type, Tg = TgF344-AD, HFHS = High-fat high-sugar diet*, ^*\**^ *= p <0.05*

Female TgF344-AD rats had comparable AUC to their WT counterparts from 0-30 min (F_1,10.21_ = 1.75, p =.215, 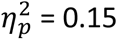; Figure 4C), 30-60 min (F_1,12.06_ = 2.48, p =.141, 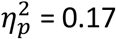; Figure 4D), 60-90 min (F_1,12.82_ = 0.001, p =.973, 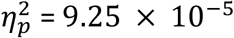; Figure 4E), and 90-120 min (F_1,12.82_ = 0.18, p =.675, 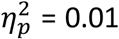; Figure 4F). There was no effect of genotype (F_1,146.02_ = 0.417, p =.519, 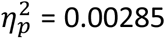) on blood glucose concentrations or on tAUC in female rats (Figure 4G). There was no significant effect of genotype on the baseline blood glucose concentrations after the 8-h fast (F_1,31.5_ = 0.1, p =.754, 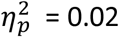; S. Figure 1D).

### 3.6. iBAT and adrenal gland mass are reduced in female TgF344-AD rats

iBAT mass was reduced in 7-month-old female TgF344-AD rats. Sex and genotype significantly interacted to impact iBAT mass (F_1,51.74_ = 12.35, p = 9.25×10-4, 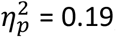; Figure 5A) such that female TgF344-AD rats had reduced iBAT mass compared to WT females (F_1,26.87_ = 13.59, p =.001, 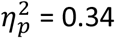). The decline was specific to females as iBAT depots in male WT and TgF344-AD rats were comparable (F_1,23.14_ = 0.88, p =.358, 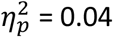).

**Figure 5:**
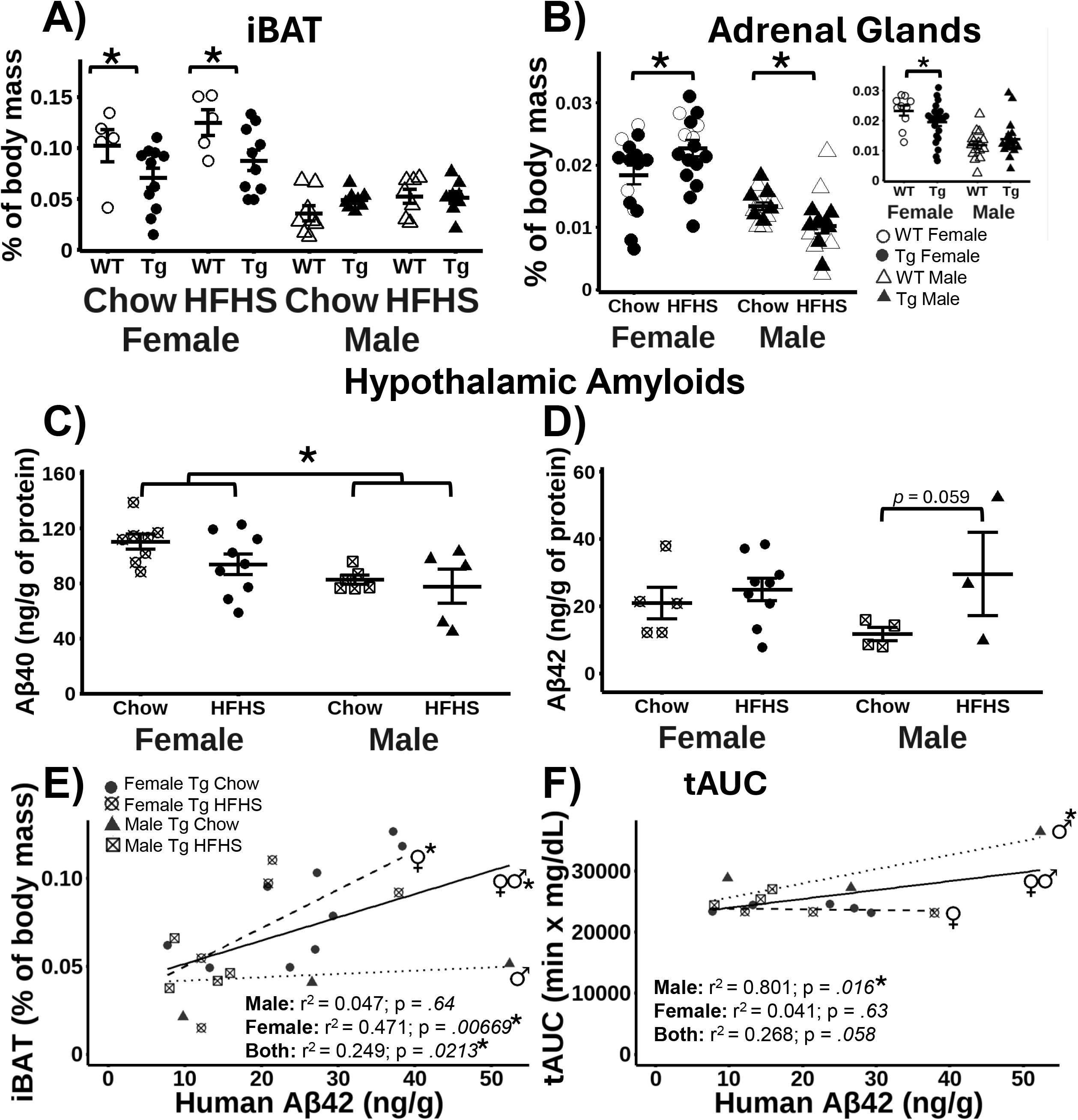
iBAT and adrenal glands mass are reduced in TgF344-AD females regardless of diet; soluble human Aβ_40_ and Aβ_42_ are expressed in the hypothalamus of prodromal TgF344-AD rats. **A)** Female TgF344-AD rats had less iBAT mass than female WT rats regardless of diet (Male: WT-Chow: n = 8, WT-HFHS: n = 7, Tg-Chow: n = 8, Tg-HFHS: n = 8; Female: WT-Chow: n = 5, WT-HFHS: n = 5, Tg-Chow: n = 11, Tg-HFHS: n = 12). **B)** Female TgF344-AD rats had reduced adrenal gland mass. HFHS diet increased adrenal mass in females and reduced it in males (Male: WT-Chow: n = 8, WT-HFHS: n = 7, Tg-Chow: n = 8, Tg-HFHS: n = 8; Female: WT-Chow: n = 5, WT-HFHS: n = 5, Tg-Chow: n = 11, Tg-HFHS: n = 12). **C)** Female TgF344-AD rats (6.8-7.3 months) had more Aβ_40_ in the hypothalamus than male rats (Male: Tg-Chow: n = 6, Tg-HFHS: n = 5; Female Tg-Chow: n = 8, Tg-HFHS: n = 9). **D)** There was a strong trend for HFHS diet-induced increases in hypothalamic Aβ_42_ in male and female Tg rats (Male: Tg-Chow: n = 4, Tg-HFHS: n = 3; Female Tg-Chow: n = 5, Tg-HFHS: n = 9). **E) i**BAT mass and hypothalamic Aβ_42_ were positively correlated in female TgF344-AD rats (Male: n = 7; Female n = 14) **F)** There was a positive association between tAUC and hypothalamic Aβ_42_ in male TgF344-AD rats. *WT = Wild-type, Tg = TgF344-AD, HFHS = High-fat high-sugar diet*, ^*\**^ *= p <0.05*

There was a significant interaction between sex and diet (F_1,52.25_ = 12.45, p = 8.75×10^−4^, 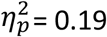 = 0.19) and between sex and genotype (F_1,54.08_ = 5.26, p =.026, 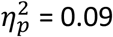) on adrenal gland mass (Figure 5B). Specifically, female TgF344-AD rats had lighter adrenal glands than their WT counterparts (F_1,27.94_ = 6.73, p =.015, 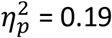), an effect that was not observed in males (F_1,21.79_ = 1.58, p =.223, 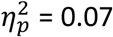). There was no impact of genotype on the mass of the heart (F_1,51.99_ = 1.02, p =.316, 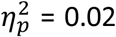; S. Figure 2B), liver (F_1,56.94_=2.08, p =.155, 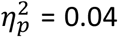; S. Figure 2C), or kidneys (F_1,50.85_ = 1.15, p =.289, 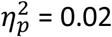; S. Figure 2D).

### 3.7. Soluble Aβ is detectable in the hypothalamus of TgF344-AD rats during the prodromal phase. Aβ_42_ correlates with iBAT mass in females and glucose dysregulation in males

Soluble human Aβ_40_ and Aβ_42_ were measured in the hypothalamus of 7-month-old TgF344-AD rats after 11 weeks on the diet (Figure 3A). In a pilot experiment, we could not detect human Aβ_40_ and Aβ_42_ in WT rats, confirming that the human Aβ assays did not cross-react with endogenous amyloid species (n = 4; data not shown). Female TgF344-AD rats had more soluble Aβ_40_ (F_1,25_=8.423, p =.008, 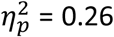; Figure 5C) than males but they did not differ in Aβ_42_ levels (F_1,25_= 0.181, p =.676, 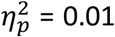; Figure 5D). The HFHS diet did not affect Aβ_40_ levels in male or female TgF344-AD rats (male: F_1,9_=0.167, p =.692, 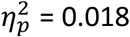; female: F_1,16_= 1.869, p =.19, 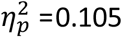 0.105). Soluble hypothalamic Aβ_42_ was undetectable in 50% of male TgF344-AD rats (4 chow, 3 HFHS) and 25% of female TgF344-AD rats (2 chow, 2 HFHS; no significant differences between sexes χ^2^ = 1.0772, df = 1, p =.2993, V = 0.189). However, there was a strong trend for the HFHS diet to increase hypothalamic Aβ_42_ in male AD rats (F_1,17_=4.106, p =.059, 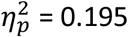; Figure 5D).

As we found evidence of iBAT dysregulation in female TgF344-AD rats and disturbed glucose regulation in male TgF344-AD rats, we sought to determine whether hypothalamic Aβ_42_ and/or Aβ_40_ correlated with iBAT mass and the AUCs during the GTT. When collapsed by sex, iBAT mass correlated positively with hypothalamic Aβ_42_ (R^2^ = 0.483, p =.0213, α_adj_ = 0.0167; Figure 5E). When separated by sex, there was a positive correlation between iBAT mass and hypothalamic Aβ_42_ in female TgF344-AD rats (R^2^ = 0.441, p = 6.69×10^−4^, α_adj_ = 0.0167), but not in male TgF-344-AD rats (R^2^ = 0.002, p =.939, α_adj_ = 0.0167). By contrast, hypothalamic Aβ_42_ correlated with AUC during the glucose tolerance test in male TgF344-AD rats (R^2^ = 0.801, p =.016, α_adj_ = 0.0167; Figure 5F), but not in female TgF344-AD rats (R^2^ = 0.041, p =.63, α_adj_ = 0.0167) or when collapsed by sex (R^2^ = 0.268, p =.058, α_adj_ = 0.0167). Hypothalamic Aβ_40_ did not correlate with iBAT, tAUC, or any of the AUC segments (data not shown).

### 3.8. TgF344-AD rats have elevated body temperatures

Body temperature was assessed between 6 and 8 months of age [45]. There was a significant interaction between genotype and light cycle for females (F_1,479_ = 8.4789, p =.00376, 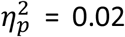), with female TgF344-AD rats exhibiting higher temperatures than female WT rats only during the dark phases (t_21.99_ = 2.49, p =.042; Figure 6B; S Figure 3A). For males, a main effect of genotype was observed (F_1,25_ = 24.31, p = 4.44×10^−5^, 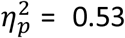), such that male TgF344-AD rats had higher body temperatures than male WT rats in both the light and dark (t_20.33_ = 3.46, p =.0049; Figure 3C; S Figure 3B). A significant interaction between genotype and light cycle (F_1,479_ = 5.112, p =.024, 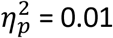) indicated that the temperature difference in the dark was greater than in the light in male TgF344-AD rats (S. Figure 3B).

**Figure 6:**
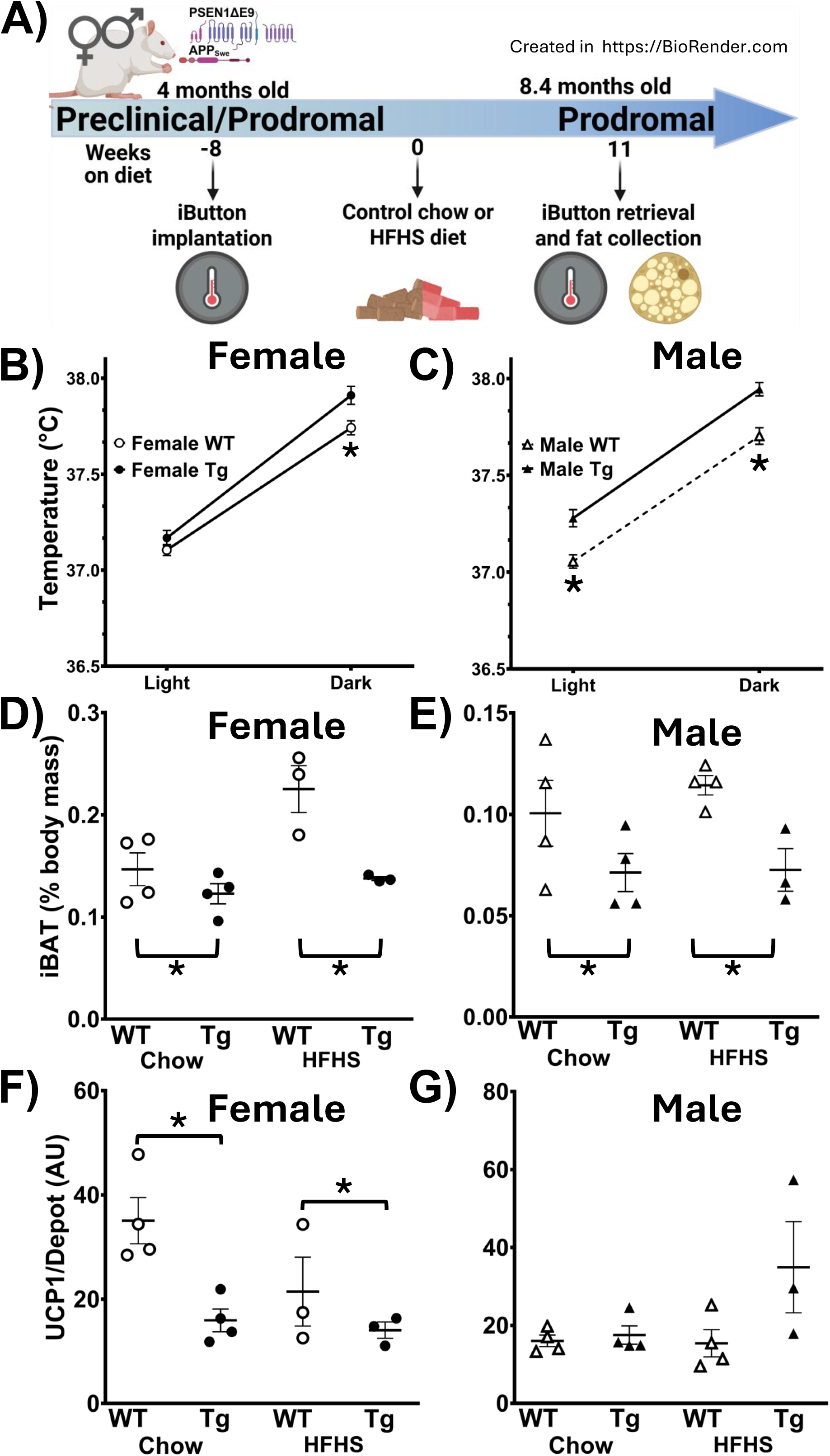
Body temperature is increased, but iBAT mass and UCP1 levels are reduced in female TgF344-AD rats. **A)** Experimental timeline. **B)** Female TgF344-AD rats (5-7 months of age) had higher body temperatures than WT females during the dark phase of the light cycle (WT: n = 13, Tg: n = 13). **C)** Male TgF344-AD rats had higher body temperatures than WT males during both the light and dark phase. (WT: n = 15, Tg: n = 14). **D)** iBAT mass was reduced in female TgF344-AD rats, (WT-Chow: n = 4, WT-HFHS: n = 4, Tg-Chow: n = 3, Tg HFHS: n = 3) and **E)** male TgF344-AD rats (WT-Chow: n = 4, WT-HFHS: n = 4, Tg-Chow: n = 4, Tg HFHS: n = 3). **F)** Female TgF344-AD rats expressed less UCP1 in iBAT tissue than WT females (WT-Chow: n =4, WT-HFHS: n = 4, Tg-Chow: n = 3, Tg HFHS: n = 3) while **G)** Male TgF344-AD rats expressed UCP1 levels comparable to WT rats (WT-Chow: n =4, WT-HFHS: n = 4, Tg-Chow: n = 4, Tg HFHS: n = 3). *WT = Wild-type, Tg = TgF344-AD, HFHS = High-fat high-sugar diet*, ^*\**^ *= p <0.05*

### 3.9. TgF344-AD rats have reduced iBAT mass, female TgF344-AD rats also have reduced UCP1

At 8.4 months of age, female and male TgF344-AD rats had reduced iBAT mass (F_1,21_ = 24.72, p =.000064, 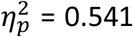; Figure 6D, E) compared to their WT counterparts. There was a significant interaction between genotype and sex for UCP1 expression (F_1,20_ = 11.18, p =.0032, 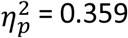; Figure 6F, G). In females, a main effect of genotype was observed (F_1,10_=10.57, p =.0087, 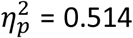), such that TgF344-AD females on both diets exhibited reduced UCP1 expression (t = 3.252, p =.0087), whereas in males, there was a trend towards increased UCP1 in TgF344-AD rats (F_1,11_= 4.227, p =.064, 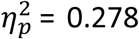). A negative association between terminal body mass and UCP1 expression was found in female rats (r^2^ = 0.336; *p* =.*0299*; α_adj_ = 0.025; S Figure 3C), and a positive association between terminal body mass and UCP1 expression was found in male rats (r^2^ = 0.287; *p* =.*0486;* α_adj_ = 0.025; S Figure 3C).

## 4. DISCUSSION

Collectively, the results of the present study show that energy homeostasis is disrupted in TgF344-AD rats and that the timing and profile of these changes depend on sex. Female, but not male TgF344-AD rats display decreases in iBAT UCP1 levels, increased energy intake and body temperature during the dark phase, and HFHS diet-induced increases in body mass. Female TgF344-AD rats also exhibit decreases in iBAT mass at an earlier age than male TgF344-AD rats. Male TgF344-AD rats have increases in body temperature during both the light and dark phase and only male TgF344-AD rats had HFHS diet-induced impairment of glucose clearance. Further, male TgF344-AD rats begin to outweigh their WT littermates during adolescence, at a younger age than female TgF344-AD rats. These increases in body mass persist for most of the prodromal period, particularly in female rats.

Our findings mechanistically add to the growing evidence indicating that energy homeostasis is disrupted in TgF344-AD rats in a sex-dependent manner. For example, older female TgF344-AD rats who are in the early clinical period (12-months-old) have more total cholesterol, free fatty acids, and high-density lipids in their blood than WT rats, whereas male TgF344-AD rats have higher triglycerides than WT rats. As in the present study, prior research has shown that female, but not male, TgF344-AD rats gain more weight than their respective WT controls on a high calorie diet (Almanza et al., 2024), and male TgF344-AD rats fed a control diet weigh more than WT rats but have comparable adiposity [51,52]. We also observed comparable adiposity between WT and TgF344-AD rats but extend those results by showing a decrease in the lean/fat ratio for TgF344-AD rats fed the control diet. Floor effects may account for the fact that a similar decrease in the lean/fat ratio was not observed in TgF344-AD rats fed the HFHS diet. Our findings also raise the possibility that increased energy intake and decreased UCP1-mediated thermogenesis contribute to these increases in weight gain in female TgF344-AD rats. This likelihood is supported by the negative correlation between iBAT UCP1 levels and body weight in female rats that we observed. A recent report showed that female, but not male, TgF344-AD rats fed a control diet had impaired glucose homeostasis at 9 and 12 months of age [52]. By contrast, we found here that there were no deficits in glucose regulation in 7-month-old chow-fed TgF344-AD rats of either sex, suggesting that our chow-fed TgF344-AD rats had not yet reached the age at which glucose intolerance develops. When fed a HFHS-diet, male TgF344-AD rats have impaired glucose regulation, suggesting that male TgF344-AD rats are more vulnerable to HFHS-induced risk of type II diabetes risk. This is consistent with findings from humans showing that males are more susceptible to type II diabetes than females, especially when young [53]. Some related sex differences have also been identified in the 3xTg-AD mouse, where 3xTgAD males but not females display increased hypothalamic inflammation when fed a control diet. When 3xTgAD mice are fed a high-fat diet, males exhibit increased systemic inflammation and elevated expression of plasma markers of diabetes [54,55]. However, in contrast to the present findings, only female 3xTg-AD mice have high fat diet-induced increases in visceral adiposity and impaired glucose clearance [55]. These differences may be because 3xTg-AD mice express human tau harboring mutations associated with frontotemporal dementia that may worsen or alter the trajectory and presentation of pathology compared to models that only express familial AD mutations [56].

The present findings appear to be the first to show that soluble Aβ is present in the hypothalamus during the prodromal period in TgF344-AD rats. This finding is consistent with observations in AD patients showing that the hypothalamus displays amyloid plaques and undergoes neurodegeneration and atrophy during early AD [13,44,57], and with the finding that glucose metabolism is reduced in the hypothalamus of presymptomatic Tg2576 mice that also overexpress APP [58]. Our data also show that there is more soluble Aβ_40_ in the hypothalamus of female than male rats, although the consequences of these elevations are unclear. Soluble Aβ_40_ reduces the deposition of soluble Aβ_42_ into plaques and protects against premature death in Tg2576 mice [59,60] but conversely may contribute to vascular pathology and its deposition co-occurs with oligomeric Aβ_42_ [61]. Interestingly, there was a trend suggesting that the HFHS diet increases hypothalamic soluble Aβ_42_ levels in male TgF344-AD rats. There were also significant sex-specific positive correlations between hypothalamic Aβ_42_ and glucose dysregulation in TgF344-AD male rats and with increased iBAT mass in TgF344-AD females.

Taken together, our data suggest that soluble human amyloid may induce hypothalamic neuropathology and produce some of the sex-specific metabolic disruptions observed in TgF344-AD rats. The paraventricular nucleus of the hypothalamus (PVN) is a major source of control over sympathetic outflow to BAT and projects to the lateral hypothalamus and the arcuate nucleus, which are critical for the regulation of feeding behavior, BAT thermogenesis, and blood glucose regulation [62,63]. Separate populations of neurons in the PVN mediate BAT thermogenesis vs. peripheral glucose homeostasis [64,65]. Since the measures of soluble Aβ in the present study were obtained from the entire hypothalamus, additional research is needed to determine whether these sex differences in metabolic disturbances are associated with differences in the hypothalamic nuclei and specific neuronal subpopulations in female vs. male TgF344-AD rats.

The present finding that adrenal gland mass is decreased in female TgF344-AD rats is inconsistent with evidence indicating that the hypothalamic-pituitary-adrenal (HPA) axis is overactivated in AD [66], given that HPA overactivation induces adrenal hyperplasia [67]. However, adrenal function has not been well-studied during the transition from MCI to AD. It is therefore possible that preclinical or prodromal AD patients might display secondary or tertiary adrenal insufficiency before going on to develop HPA overactivation in later stages. Secondary and tertiary adrenal insufficiency are due to pituitary or hypothalamic dysfunction, respectively, which could reduce adrenal gland size [68].

The finding that TgF344-AD rats have higher body temperature than WT rats is novel, intriguing, and consistent with evidence showing that AD patients have increased core body temperature [69]. The increases we observed are not likely due to elevated activity levels, because home cage activity and locomotor movement and speed are not increased in male or female TgF344-AD rats across a wide range of ages and measures [24,25,45,70,71]. The elevations in body temperature during the dark phase observed in female TgF344-AD rats are surprising because UCP1 levels were reduced in these rats. We speculate that this increase that is specifically during the dark is due to their elevated energy intake that also occurred only during the dark and the associated increase in activity and digestion-induced thermogenesis. Given that male TgF344-AD rats did not consume more food, did not have significant reductions in UCP1, and that UCP1 was positively associated with terminal body mass, the processes that contributed to their elevated body temperatures remain unclear. One possibility is that UCP1-independent mechanisms of thermogenesis mediate these increases in male TgF344-AD rats, such as UCP3-related non-shivering thermogenesis in skeletal muscle [72].

Our novel findings that iBAT hypertrophy correlates with hypothalamic Aβ_42_ in female rats suggests that hypothalamic dysfunction may mediate the iBAT disturbances. As female TgF344-AD rats also exhibited reduced iBAT UCP1 expression, this association may be evidence of the whitening of the iBAT and its accumulation of excess lipids [73,74]. Elevations in body temperature and iBAT dysfunction have also been documented in transgenic mouse models of AD such as the 3xTgAD and APP/PS1 mice [75-77]. Exogenous and transgenic increases in Aβ_42_ promote excitability in NPY-expressing cells in the arcuate nucleus [78], which would be expected to reduce iBAT-mediated thermogenesis [79]. Further, elevations in glucose inhibit arcuate nucleus NPY neurons; as a result, Aβ_42_-induced hyperexcitability of NPY neurons would be expected to impair their detection of hyperglycemia [80,81]. Relatedly, intracerebroventricular infusions of Aβ oligomers induce hypothalamic inflammation that precedes glucose intolerance [82]. Collectively, these lines of evidence suggest that soluble Aβ oligomers may drive hypothalamic excitability to induce peripheral metabolic dysfunction in a sex-specific manner. Further research is necessary to confirm that hypothalamic Aβ mediates the sex-specific changes in iBAT and glucose regulation in TgF344-AD rats. The coincidence of peripheral metabolic dysfunction with AD is well established, but the presentation of specific metabolic dysfunctions is heterogenous in the patient population [83,84]. Our findings suggest that elevated soluble amyloid-β may drive some of this heterogeneity by interacting with biological sex.

Our findings raise the intriguing possibility that metabolic disturbances are an early symptom of AD. In humans, AD development typically starts in midlife, and metabolic disturbances during this period, such as obesity, are associated with increased AD risk [85,86]. We and others have shown that obesity exacerbates AD-Like pathology [45,55,87]. For example, we recently showed that feeding TgF344-AD rats the HFHS diet during the early prodromal period negatively impacts AD trajectory [45]. The negative effects include the induction of memory deficits in TgF344-AD rats that are not observed in chow-fed TgF344-AD rats or in HFHS-fed WT rats, increased expression of genes suggestive of mitochondrial dysfunction and lipid accumulation in the dorsal hippocampus (dHC), and disrupted dHC norepinephrine turnover. Collectively, that evidence suggests that obesity and associated comorbid conditions during early AD increases AD risk. The present findings suggest that obesity during midlife is a risk factor for AD, at least in part, because it is an early symptom of AD.

In summary, we show that energy homeostasis is disrupted in TgF344-AD rats during early AD development in a sex-dependent manner, and that some of these changes are associated with hypothalamic Aβ_42_ levels. The presence of sex-dependent associations between hypothalamic Aβ_42_ levels and measures of disrupted energy homeostasis highlight the need for future research into hypothalamic dysfunction in AD with mixed-sex study designs and raise the possibility that the occurrence of early disruptions of energy homeostasis may serve as biomarkers of AD or increased AD-risk.

## 5. DATA AVAILABILITY

Data supporting the findings can be found online at Mendeley’s Data Repository with the DOI: 10.17632/rthpp2ktb7.1

## 6. ACKNOWLEDGEMENTS

We thank Trenton Buckner, Kam-Ren Douglas, Kate Halfon, Harish Koyya, Hanh “Vivian” Ngo Vu, and Usama Zafar for their technical assistance, and Georgia State University Division of Animal Resources for their exceptional work and care.

## FIGURE CAPTIONS

**Supplemental Figure 1:**
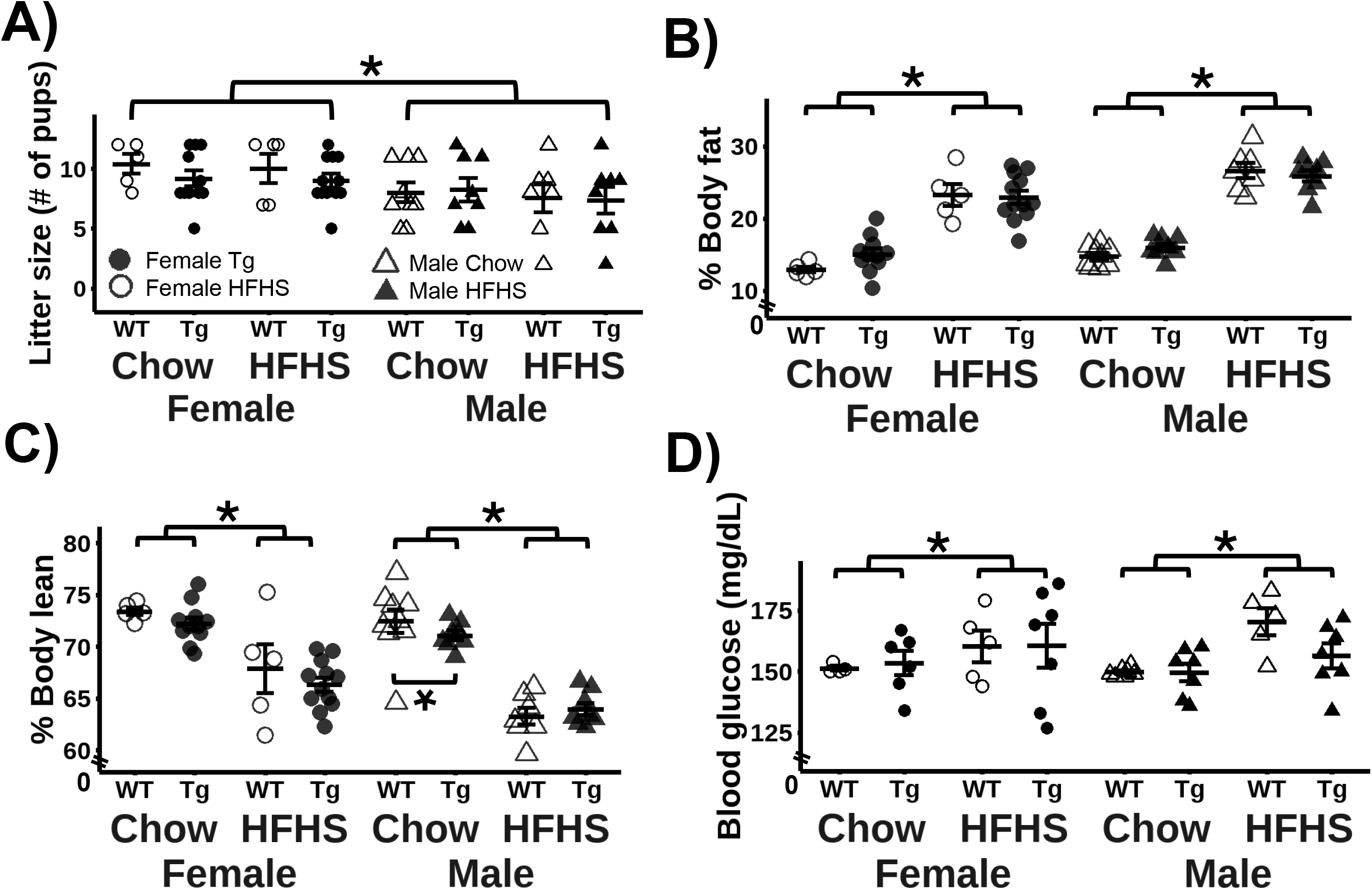
Litter size differs between male and female rats and the HFHS diet increased fasting blood glucose. **A)** Females came from larger litters than males (F_57_ = 7.795, p =.007, 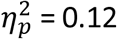), necessitating the use of litter size as a random effect in the analyses of metabolic and morphological metrics. There was no effect of genotype on litter size (F_57_ = 0.67, p =.479, 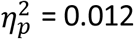). **B)** The HFHS diet increased fat mass in male and female rats (6.3-7 months) irrespective of genotype (Male: WT-Chow: n = 9, WT-HFHS: n = 7, Tg-Chow: n = 8, Tg-HFHS: n = 8; Female: WT-Chow: n = 5, WT-HFHS: n = 5, Tg-Chow: n = 11, Tg-HFHS: n = 12). **C)** The HFHS diet reduced lean mass in males and female rats irrespective of diet. Male TgF344-AD rats fed chow had less lean mass than WT-chow males (Male: WT-Chow: n = 9, WT-HFHS: n = 7, Tg-Chow: n = 8, Tg-HFHS: n = 8; Female: WT-Chow: n = 5, WT-HFHS: n = 5, Tg-Chow: n = 11, Tg-HFHS: n = 12). **D)** The HFHS diet increased fasting blood (F_38.98_ = 11.71, p =.001, 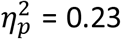) regardless of sex or genotype (Male: WT-Chow: n = 7, Tg-Chow: n = 7, WT-HFHS: n = 6, Tg-HFHS: n = 8; Female: WT-Chow: n = 4, Tg-Chow: n = 9, WT-HFHS: n = 5, Tg-HFHS: n = 10). *WT = Wild-type, Tg = TgF344-AD, HFHS = High-fat high-sugar diet*, 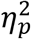 = *partial eta squared*, ^*\**^ *= p <0.05*

**Supplemental Figure 2.**
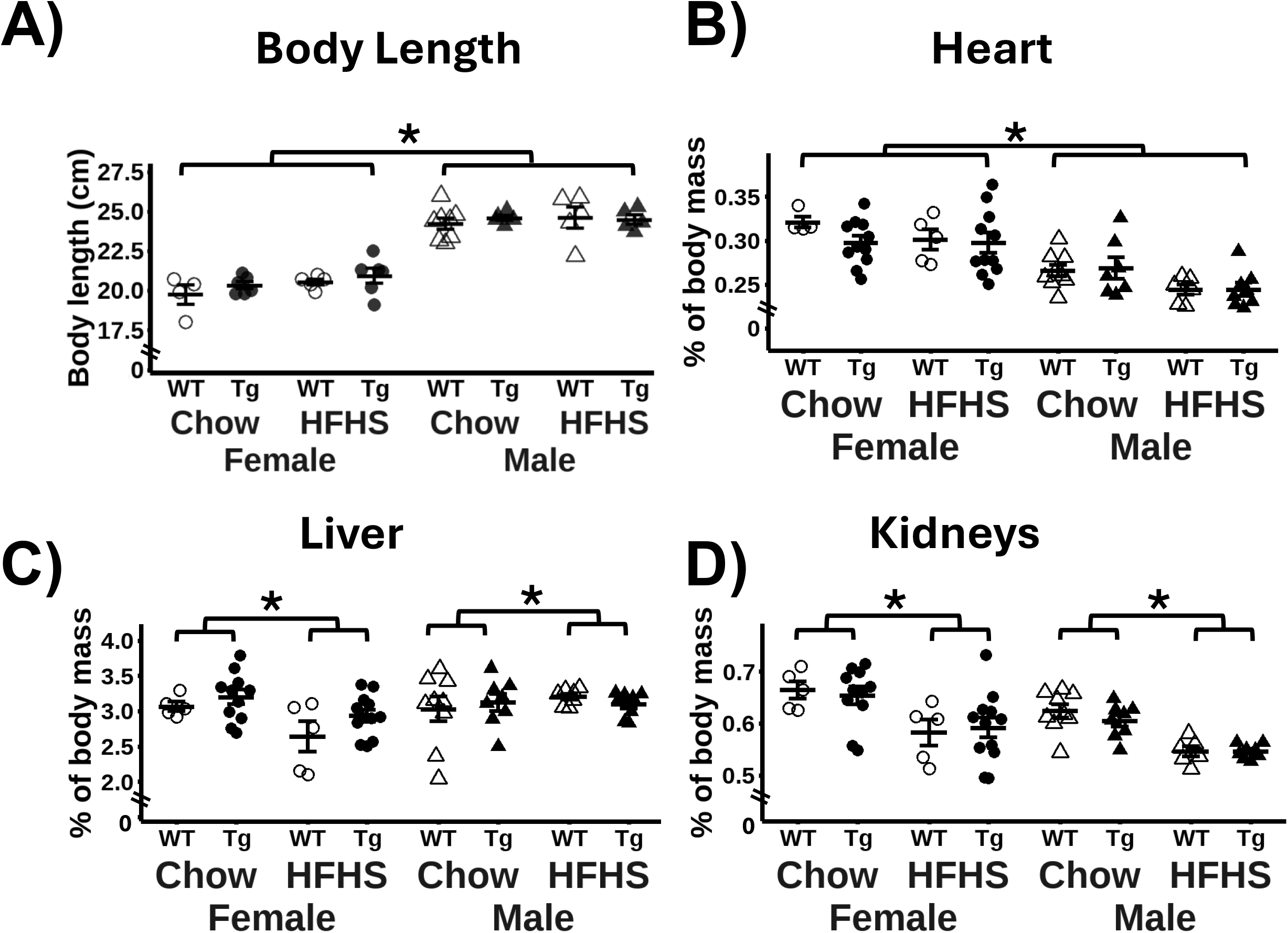
Body length, heart mass, liver mass, and kidney mass are not affected in TgF344-AD rats. **A)** There was no effect of genotype on body length (Males: WT-Chow: 8, WT-HFHS: 5, Tg-Chow: n = 5, Tg-HFHS: n = 5; Females: WT-Chow: n = 4, WT-HFHS: n = 5, Tg-Chow: n = 6, Tg-HFHS: n = 6). **B)** Females have smaller hearts than males (F_1,42.2_ = 38.6, p = 1.93×10^−7^, 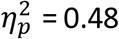;Male: WT-Chow: n = 9, Tg-Chow: n = 7, WT-HFHS: n = 7, Tg-HFHS: n = 7; Female: WT-Chow: n = 5, Tg-Chow: n = 11, WT-HFHS: n = 5, Tg-HFHS: n = 11). **C)** The HFHS diet reduced liver mass in female rats (Male: F_1,23.06_ = 0.07, p =.796, 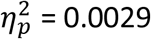; Female F_1,28.18_ = 22.17, p = 1.77×10^−5^, 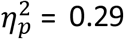; Male: WT-Chow: n = 9, Tg-Chow: n = 8, WT-HFHS: n = 8, Tg-HFHS: n = 7; Female: WT-Chow: n = 5, Tg-Chow: n = 11, WT-HFHS: n = 5, Tg-HFHS: n = 12). **D)** The HFHS diet reduced kidney mass in male and female rats (Male: F_1,14.65_ = 32.51, p = 4.59×10^−5^, 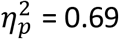; Female: F_1,28.21_ = 10.73, p =.003, 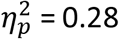; Male: WT-Chow: n = 8, Tg-Chow: n = 8, WT-HFHS: n = 7, Tg-HFHS: n = 7; Female: WT-Chow: n = 5, Tg-Chow: n = 11, WT-HFHS: n = 5, Tg-HFHS: n = 11*). WT = Wild-type, Tg = TgF344-AD, HFHS = High-fat high-sugar diet*, 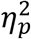 *= partial eta-squared*, ^*\**^ *= p <0.05*

**Supplemental Figure 3:**
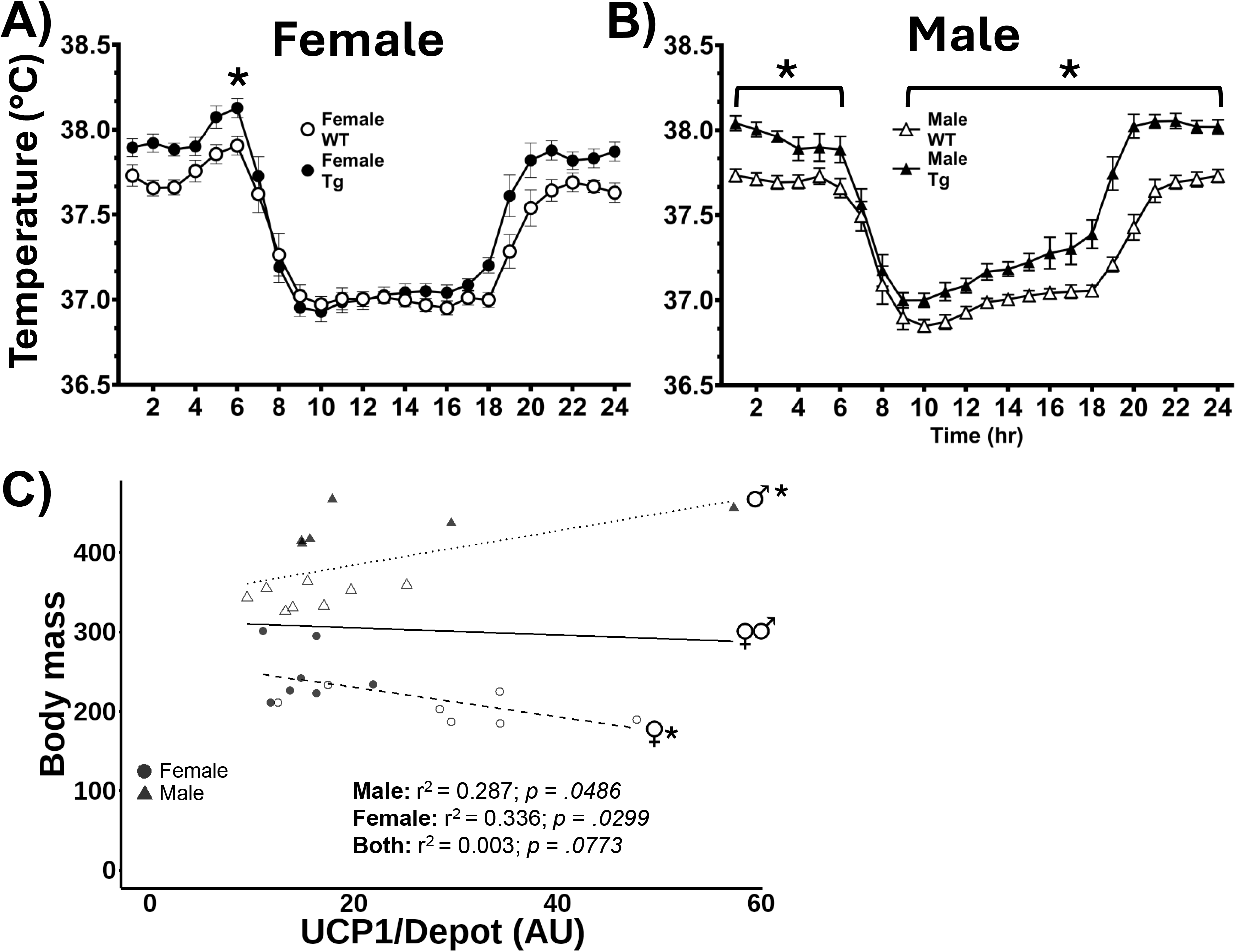
Male TgF344-AD rats have higher body temperatures than WT males regardless of diet or time of day; female TgF344-AD rats have higher body temperatures only during the dark phase. **A)** In female rats, there was an interaction between hour and genotype (F_23,437_ = 1.838, p =.0109, 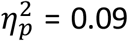). TgF344-AD females had higher temperatures at hour 2 of the dark phase (t = 3.69, df = 19, p =.036). **B)** In male rats, there was an interaction between hour and genotype (F_23,391_ = 1.659, p =.0297, = 0.09) such that male TgF344-AD rats had higher body temperatures than WT rats at all timepoints (t: 3.78-7.301, df = 17, p:.0358-.0000297) except hours 6,7, and 8. **C)** A negative association between body mass and iBAT UCP1 expression was found in female rats (p =.0299, n = 14) and a positive association was found in male rats (p =.0486, n = 15). *WT = Wild-type, Tg = TgF344-AD, HFHS = High-fat high-sugar diet*, 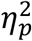 *= partial eta squared*, ^*\**^ *= p <0.05*

**Supplemental Table 1.** Caloric content of the standard chow diet and HFHS diet.

| **Macronutrients**<br>**(% kcal)** | **Lab Diet**<br>**5001** | **Research Diets**<br>**D12451 (HFHS)** |
| --- | --- | --- |
| **Protein** | **28.9** | **20** |
| **Fat** | **13.6** | **45** |
| **Carbohydrate** | **57.5** | **35** |
| **Sucrose** | **3.25** | **17** |
| **kcal/g** | **3.36** | **4.73** |

**Supplemental Table 2.** Number of rats analyzed for each measure and reasons for any exclusions.

|  | Experimental outliers |  |  | Statistical outliers |  |  |  |  |  |  |  |
| --- | --- | --- | --- | --- | --- | --- | --- | --- | --- | --- | --- |
|  |  |  | Total<br>Remaining | Female |  |  |  | Male |  |  |  |
|  | Total<br>Tested | Exclusion reason |  | WT-Chow | WT-HFHS | Tg-Chow | Tg-HFHS | WT-Chow | WT-HFHS | Tg-Chow | Tg-HFHS |
| Experiment 1 |  |  |  |  |  |  |  |  |  |  |  |
| Body mass- cross sectional | 75 |  | 75 | 1 | NA | 0 | NA | 0 | NA | 0 | NA |
| Early body mass- longitudinal | 127 |  | 127 | 8 | NA | 3 | NA | 2 | NA | 3 | NA |
| Caloric intake- food consumption | 31 | food hopper error | 29 | 0 | 0 | 0 | 0 | 2 | 2 | 0 | 0 |
| Experiment 2 |  |  |  |  |  |  |  |  |  |  |  |
| Body temperature | 56 |  | 56 | 0 | 0 | 0 | 1 | 1 | 0 | 0 | 0 |
| iBAT mass- body mass | 31 | Collection error | 30 | 0 | 0 | 0 | 0 | 0 | 0 | 0 | 1 |
| iBAT- UCP1 per depot | 30 | Collection error | 29 | 0 | 0 | 0 | 0 | 0 | 0 | 0 | 0 |
| Experiment 3 |  |  |  |  |  |  |  |  |  |  |  |
| Body mass- longitudinal | 65 |  | 65 | 0 | 0 | 0 | 1 | 0 | 0 | 1 | 0 |
| Body mass- terminal | 65 |  | 65 | 0 | 0 | 1 | 1 | 0 | 0 | 0 | 0 |
| Body length | 65 | Collection error | 45 | 0 | 0 | 0 | 0 | 0 | 0 | 1 | 0 |
| Blood glucose - GTT | 49 |  | 49 | 1 | 0 | 2 | 2 | 2 | 1 | 1 | 0 |
| GTT AUC | 49 |  | 49 | 1 | 0 | 3 | 2 | 2 | 1 | 1 | 0 |
| Blood glucose - fasting | 65 |  | 49 | 0 | 0 | 0 | 0 | 1 | 0 | 0 | 0 |
| EchoMRI- Fat mass | 65 |  | 65 | 0 | 0 | 0 | 0 | 0 | 0 | 0 | 0 |
| EchoMRI- Lean mass | 65 |  | 65 | 0 | 0 | 0 | 0 | 0 | 0 | 0 | 0 |
| igWAT- mass | 65 |  | 65 | 0 | 0 | 0 | 0 | 0 | 0 | 1 | 0 |
| rpWAT- mass | 65 |  | 65 | 0 | 0 | 2 | 2 | 0 | 0 | 0 | 0 |
| gWAT- mass | 65 |  | 65 | 0 | 0 | 0 | 0 | 0 | 0 | 0 | 0 |
| iBAT- mass | 65 | Collection error | 63 | 0 | 0 | 0 | 1 | 0 | 0 | 0 | 0 |
| Adrenal glands- mass | 65 |  | 65 | 0 | 0 | 0 | 0 | 1 | 0 | 0 | 0 |
| Heart- mass | 65 | Collection error | 64 | 1 | 0 | 0 | 0 | 0 | 1 | 1 | 0 |
| Liver- mass | 65 |  | 65 | 0 | 0 | 0 | 0 | 0 | 0 | 0 | 0 |
| Kidney- mass | 65 |  | 65 | 0 | 1 | 0 | 0 | 0 | 0 | 0 | 0 |
| Hypothalamic A$\beta_{40}$ | 32 | Low CV | 28 | NA | NA | 0 | 0 | NA | NA | 0 | 0 |
| Hypothalamic A$\beta_{42}$ | 32 | Undetectable | 21 | NA | NA | 0 | 0 | NA | NA | 0 | 0 |

